# Region-specific variations in the cerebrovasculature underlie disease progression in Parkinson’s disease

**DOI:** 10.1101/2025.03.09.641541

**Authors:** Derya Dik, Glenda M. Halliday, Vladimir Sytnyk, Claire E. Shepherd

## Abstract

Parkinson’s disease is a progressive neurodegenerative disorder characterised by motor dysfunction, dopaminergic neuronal loss in the substantia nigra and abnormal accumulation of α-synuclein Lewy bodies. Research suggests that the cerebrovascular system plays a role in fluid dynamics, waste clearance, and removal of abnormal proteins. Imaging studies show that this waste clearance system, known as the glymphatic system, is disrupted in Parkinson’s disease, highlighting its involvement in the disease.

This immunohistochemical human brain tissue study quantified changes in the cerebrovascular system (perivascular space, string vessels, pericytes, aquaporin-4 and astrocytes) in Parkinson’s disease (n=18) cases with variable disease durations (median=14, range= 19) compared to age and post-mortem matched (*P* >0.05) control cases (n=7). Analysis was carried out in brain regions variably affected by cell loss (substantia nigra) and protein deposition (substantia nigra and medial temporal cortex). The occipital cortex was included, as this region is not affected by cell loss or protein deposition. Group differences were analysed and the relationship with protein deposition (Lewy body stage, amyloid score, neurofibrillary tangle score) was assessed.

Although total astrocyte density did not change (*P* >0.05), Parkinson’s disease cases exhibited reduced aquaporin-4 in astrocytic endfeet and enlargement of the arteriolar and venular perivascular space. Significant changes in the capillary network were also observed with increased string vessel formation (*P* <0.001) and pericyte loss (*P* <0.001), changes likely to impact blood flow and its regulation. The formation of string vessels significantly correlated with disease duration (*P* <0.05), especially in the occipital cortex. The occipital cortex demonstrated the greatest decreases in pericytes (*P* <0.001) and aquaporin-4 mislocalisation (*P* <0.05), while changes in pericyte density were also significant in the substantia nigra. In contrast, these changes were not significant in the medial temporal cortex despite protein deposition in this region. Although no Lewy pathology was detected in the occipital cortex, there was a positive relationship between Lewy body stage and perivascular space size (Rho =0.6, *P* <0.05).

These findings reveal progressive, region-specific alterations in the cellular components of the glymphatic system and vascular integrity in Parkinson’s disease. Notably, the correlation between string vessel formation and disease duration, even in a region unaffected by protein deposition, suggests that vascular changes may play an important role in disease progression. These results emphasize the need for further investigation into the interplay between regional vascular changes and Parkinson’s disease progression, which may offer novel insights for therapeutic strategies.

## Introduction

Parkinson’s disease (PD) is a progressive neurodegenerative disorder characterised by motor symptoms including bradykinesia, tremor and rigidity.^1^ The underlying cause of idiopathic PD is still unknown although the loss of dopaminergic neurons in the substantia nigra and brainstem α-synuclein deposition in the form of Lewy bodies (LB) and Lewy neurites (LN) are diagnostic.^2^ Lewy pathology spreads progressively in PD in a characteristic pattern throughout brainstem (stage I-IV), limbic (stage V) and neocortical (stage VI) brain regions.^3,4^ Non-motor symptoms such as visual hallucinations and fluctuating cognition can also be present in advanced stages of PD,^5^ or in individuals with a later age of disease onset.^6^ Later age of disease onset is also associated with greater burdens of other pathologies, including β-amyloid plaques and neocortical neurofibrillary tangles (NFT).^7^

Recent studies have identified the ‘glymphatic system’ comprising a network of perivascular spaces enclosed by the endfeet of astrocytes that supports the clearance of waste proteins from the brain.^8–10^ This system shows increased activity during sleep,^11^ and is impaired by sleep disruption,^12^ which is a common and early feature of PD.^13^ Indeed, diffusion tensor image analysis along the perivascular space (DTI-ALPS) has identified alterations in fluid flow in PD in regions with significant α-synuclein deposition.^14^ Correlations between glymphatic dysfunction and clinical symptoms such as motor and cognitive impairment have also been noted.^15,16^ Animal models of PD and glymphatic flow reveal that blocking meningeal lymphatic vessels in mice with the human *SNCA* A53T gene mutation led to α-synuclein deposition and motor dysfunction,^17^ thereby providing proof of concept that this system may play an important role in the deposition of α-synuclein. Polarisation of the aquaporin-4 (AQP4) water channel to the endfeet of astrocytes also plays an essential role in this system, with animal studies showing a 70-80% reduction in interstitial solute clearance in AQP4-deficient mice.^9^

Brain imaging studies have identified white matter hyperintensities, an indicator of small vessel disease, in individuals meeting the clinical criteria for PD, particularly those with cognitive decline.^18–20^ In contrast, post-mortem studies of neuropathologically confirmed PD cases have consistently reported low levels of cerebrovascular changes that would be detected through imaging.^21^ Interestingly, some of these studies suggest an inverse relationship between vascular risk factors, cerebrovascular disease, and PD.^22,23^ More recently, human brain tissue studies have identified a decrease in AQP4 in astrocytic endfeet in the frontal cortex of PD patients.^24^ This finding suggests potential disruptions in fluid transport across the blood-brain barrier (BBB) in PD. Furthermore, alterations in the capillary network and endothelial cell integrity have also been observed in PD brain.^25–28^

This study examined changes in the perivascular space (PVS), AQP4 localisation, astrogliosis, capillary network and pericyte number in PD cases with differing disease durations, compared to age-matched controls. Our analysis focused on brain regions differentially affected by protein deposition and cell loss. We hypothesise that disruptions in the cellular components of the glymphatic system exacerbate the accumulation of pathological proteins such as α-synuclein, and contribute to disease progression in PD.

## Materials and methods

### Brain tissue samples

Tissue samples from pathologically confirmed and staged cases of PD (n=18)^4^ and controls without neurological or neuropathological disease (n=7) were obtained from the Sydney Brain Bank following project-specific ethics approval (University of New South Wales Human Research Ethics Advisory Committee HC220172). All cases with PD were levodopa-responsive and fulfilled the UK Brain Bank Clinical Criteria for a diagnosis of clinical PD^29^ with no other neurodegenerative conditions (see A and B score, **Table 1**), neoplasms, or significant infarctions. The details for the groups are given in **Table 1**.

**Table 1.**
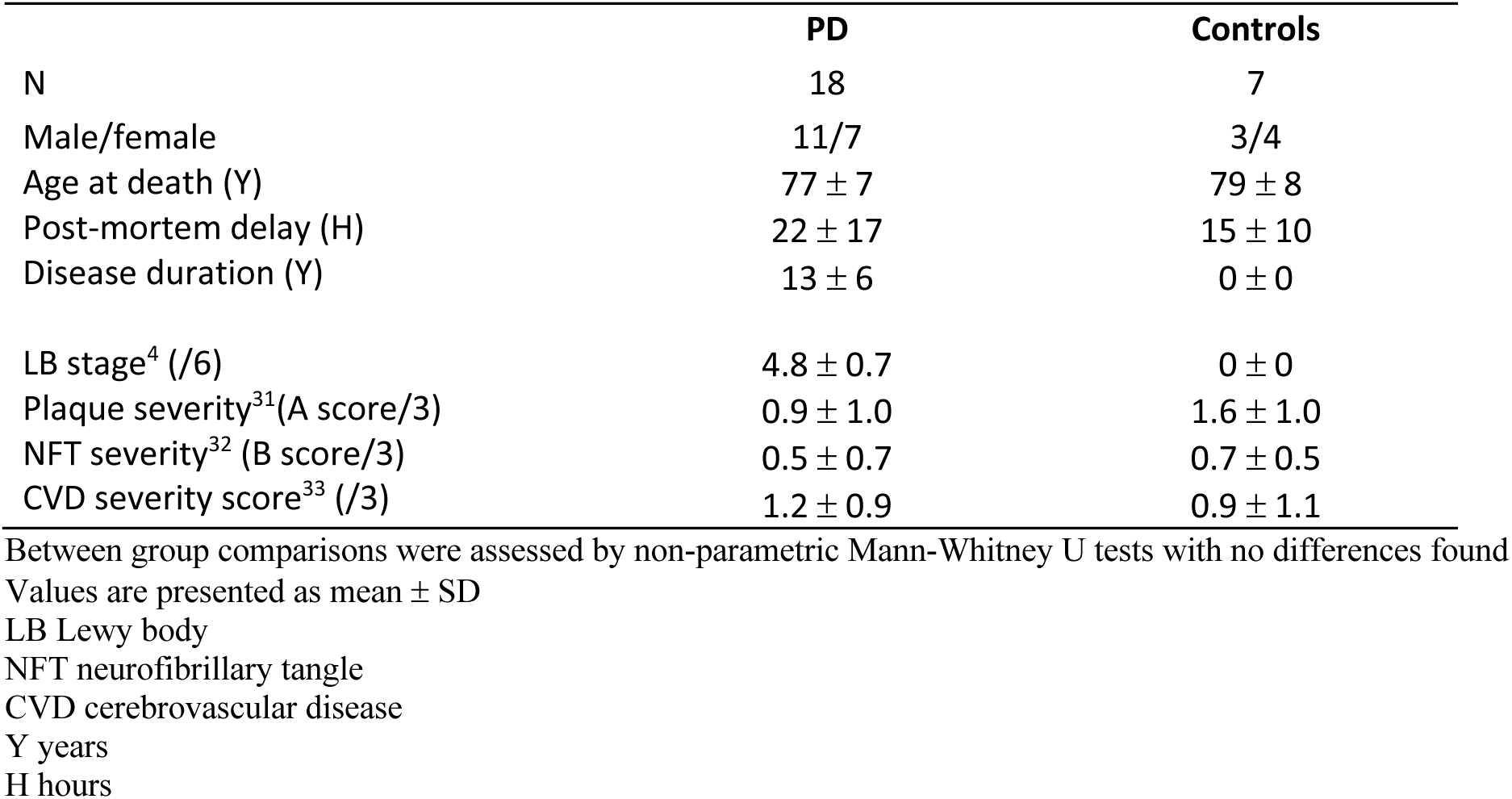
Demographic data for the PD and the control groups.

|  | PD | Controls |
| --- | --- | --- |
| N | 18 | 7 |
| Male/female | 11/7 | 3/4 |
| Age at death (Y) | 77 ± 7 | 79 ± 8 |
| Post-mortem delay (H) | 22 ± 17 | 15 ± 10 |
| Disease duration (Y) | 13 ± 6 | 0 ± 0 |
| LB stage<sup>4</sup> (/6) | 4.8 ± 0.7 | 0 ± 0 |
| Plaque severity<sup>31</sup> (A score/3) | 0.9 ± 1.0 | 1.6 ± 1.0 |
| NFT severity<sup>32</sup> (B score/3) | 0.5 ± 0.7 | 0.7 ± 0.5 |
| CVD severity score<sup>33</sup> (/3) | 1.2 ± 0.9 | 0.9 ± 1.1 |

Between group comparisons were assessed by non-parametric Mann-Whitney U tests with no differences found Values are presented as mean ± SD
LB Lewy body
NFT neurofibrillary tangle
CVD cerebrovascular disease
Y years
H hours

Three mm thick coronal formalin-fixed tissue blocks were sampled from the substantia nigra at the level of the exiting third nerve, medial temporal cortex including the hippocampus and entorhinal cortex at the level of the lateral geniculate nucleus and occipital striate cortex. These regions were chosen because of their varying levels of cell loss (substantia nigra) and pathology (substantia nigra and medial temporal cortex), while the occipital cortex was selected for comparison because it is not affected by either. Blocks were embedded in paraffin and twelve serial 10μm thick sections were cut and dried onto slides. Serial sections were required for this study to ensure that structures could be examined in close proximity while analysing several cellular components using different immunohistochemical techniques. Sections were deparaffinised using xylene and graded alcohols, then processed using microwave antigen retrieval (sections boiled for three minutes in 0.1M citrate buffer) for staining, as described below. Inferior temporal cortex tissue sections from an individual with mixed neuropathologies were used as both positive and negative controls for each stain. For the negative controls, the primary antibody was omitted to confirm there was no non-specific binding of the secondary antibodies. No immunoreactivity was observed in these negative control sections. **Supplementary Table 2** provides detailed information on the sections stained and their corresponding staining.

### Immunofluorescence

Section 5 of the serial sequence was used for analysis of the PVS using antibodies to detect vascular endothelial cells (using rabbit anti-Factor VIII), smooth muscle (using mouse anti-SMA-Cy3) and astrocytes (using goat anti-GFAP). Smooth muscle actin was used to differentiate between arterioles and venules to assess potential differences in PVS between these vessel types. Tissue sections were incubated at 37°C for two hours, washed, and then incubated overnight at 37°C in the dark with secondary fluorophores corresponding to anti-Factor VIII and anti-GFAP. SMA was conjugated to Cy3 and hence a secondary detection antibody was not required. Autofluorescence was quenched by TrueBlack in 70% ethanol for five minutes, and the sections washed thoroughly prior to incubating with DAPI solution for at least one and a half hours in the dark at room temperature. Sections were then washed and coverslipped with an aqueous mounting medium.

Section 8 was used for immunofluorescent staining to examine the capillary network and pericytes (using goat anti-collagen IV and DAPI counterstain, **supplementary Table 1 for antibody details**). Section 6 was used for immunofluorescence to assess AQP4 localisation to astrocyte endfeet (using rabbit anti-AQP4 and DAPI counterstain, **supplementary Table 1 for antibody details**). Tissue sections were incubated overnight at 4°C, washed and then incubated with their corresponding secondary fluorophores in the dark for one hour at room temperature. Autofluorescence was quenched, the sections washed thoroughly prior to incubating with DAPI solution and coverslipping (see above).

### Peroxidase immunohistochemistry

Peroxidase immunohistochemistry was performed using manual and automated techniques. Prior to use, tissue sections were pre-treated with 1% H_2_O_2_ in 50% ethanol for 30 minutes and 10% normal horse serum for 20 minutes at room temperature. Serial sections (**supplementary Table 2**) were stained for either collagen IV to detect the basement membrane for the analysis of capillary networks (using goat anti-collagen IV, **supplementary Table 1**), or with an antibody cocktail to detect astrocytes (rabbit anti-GFAP, mouse anti-S100β and mouse anti-ALDH1L1, **supplementary Table 1**). The antibodies were incubated on the tissue sections overnight at 4°C. The sections were washed and incubated with their corresponding biotinylated secondary antibodies for 30 minutes at 37°C. The sections were then washed and incubated with VECTASTAIN avidin-biotin tertiary antibody complex for 30 minutes at room temperature, visualised with 3,3’-diaminobenzidine (DAB) (Vector Laboratories, SK4105), and counterstained with hematoxylin. The sections were then dehydrated through a series of graded alcohols and xylene, and coverslipped using DPX. It is important to note that sections incubated with collagen IV were not counterstained, as a clean background was required to detect the capillary network and identify string vessels.

Pre-treatment with concentrated formic acid for three minutes was required prior to staining with antibodies against β-amyloid and α-synuclein (**supplementary Table 1**). Tau was identified using an antibody against hyperphosphorylated tau (**supplementary Table 1**). Staining was undertaken using the BenchMark GX autostainer with Optiview Detection kit (760-700, Roche Diagnostics) and a haematoxylin counterstain.^30^

### Imaging of tissue sections

Images of whole tissue sections were acquired using the Olympus VS200 Research Slide Scanner. Both brightfield and immunofluorescent sections were initially scanned with the 2.5x objective for characterisation and navigation of the region. The tissue sections were then imaged with the 20x objective for analysis. Five (pericytes, capillaries, string vessels and astrocytes) or six regions of interest (PVS and AQP4) from each tissue section were randomly sampled using either QuPath (Queen’s University Belfast, version 3.2) or Fiji Image J software (version 2.14.0/1.54f), with each sample area measuring approximately 0.32mm^2^ (561μm x 561μm).

### Quantitation

#### Analysis of pathological protein deposition and cerebrovascular disease

Pathological severity scores were provided by the Sydney Brain Bank. Braak LB stage was generated according to Braak *et al*.^4^ and α-synuclein positive LNs and LBs were scored as either presence or absence in the midbrain and medial temporal regions. β-amyloid A scores (represents the spread of β-amyloid positive plaque pathology) and tau B scores (represents the spread of NFT pathology) were generated using simplified criteria.^31,32^ The severity of cerebrovascular disease (CVD) was assessed according to Esiri *et al*.^33^

In addition, a serial section taken from the occipital striate cortex **(supplementary Table 2)** was stained using α-synuclein immunohistochemistry (**supplementary Table 1**). α-synuclein pathology was scored as either presence or absence in this region.

#### Quantitation of perivascular space

The PVS was identified in triple labelled immunofluorescent sections (**supplementary Fig. 1**) stained for vascular endothelial cells (red), arteriolar smooth muscle cells (yellow) and astrocytes (green, see **supplementary Table 1** for antibody details and **supplementary Table 2** for tissue section stained). Arterioles were differentiated from venules by the presence of a prominent smooth muscle layer. Blood vessels sampled for analysis ranged in cross-section from approximately 20-60μm in size.

QuPath was used to randomly sample six blood vessels per region per case; three arterioles labelled intensely using anti-SMA and three venules that were not strongly immunoreactive against anti-SMA. Regions analysed were the medial and lateral substantia nigra, and the medial temporal and occipital striate cortical gray matter. The size of the PVS was determined using two custom Fiji plug-ins. The images were exported into Fiji and ‘objects’ required for analysis detected through pixel intensity on their corresponding fluorescent channels. For this analysis, the inner (endothelium) and outermost (astrocyte) boundary of the blood vessel were the objects of interest. Manual deselection was applied to objects not necessary for analysis (i.e. insignificant background). Two separate mask-like images were created for each vessel using the endothelial and astrocyte markers **(supplementary Fig. 1)**. A distance map was generated, where each pixel in the image was assigned a value based on the distance between the inner and outer boundary **(supplementary Fig. 1F)** and the mean distance between the two objects determined. The mean distance for venules (venular PVS) and arterioles (arteriolar PVS) was averaged per case and used for statistical analysis. Venule and arteriole measures were also averaged together to provide a combined mean distance value (total PVS).

For validation of this method (as well as the quantitative methods below), five randomly selected images from two different cases were counted by two researchers, with values varying between researchers by less than 5%, indicating high inter-rater reliability.

#### Analysis of normal capillaries and capillary string vessels

Capillaries and string vessels were identified using anti-collagen IV peroxidase immunohistochemistry (**supplementary Fig. 2A-B**), with capillaries defined by their tubular appearance within the tissue **(supplementary Fig. 2A)**, and string vessels having a string-like appearance branching off or between a defined capillary **(supplementary Fig. 2B)**.

QuPath was used to randomly sample five 561μm x 561μm areas per region per case. Regions analysed were the medial and lateral substantia nigra, and the medial temporal and occipital striate cortical gray matter. QuPath was programmed to recognise capillaries from insignificant background tissue through image training via annotation and pixel classification. Hierarchical classes were created for different objects in each of the areas sampled (i.e. tissue background and blood vessels). Training parameters such as pixel classification resolution and minimum object size were set to eliminate inconsistencies in the defined annotations (this prevented inclusion of unwanted objects in the analysis, such as background staining). QuPath training was repeated until the software could reliably identify all capillaries.

QuPath was unable to distinguish between normal capillaries and string vessels, making it necessary to manually segment normal capillaries from their branching string vessels using a QuPath annotation tool. Briefly, the ‘magic wand’ tool was used to isolate the string vessels from the normal capillaries using colour intensity (DAB peroxidase). All annotations were then exported into Fiji using a custom QuPath Groovy script to analyse total length and to determine the proportion attributable to string vessels. Total capillary length and the percentage of string vessels were averaged per case and these were the values used for statistical analysis.

#### Quantitation of pericytes

Pericytes were identified on normal capillaries using immunofluorescent sections stained for capillary basement membrane (anti-collagen IV, green) with pericytes identified by DAPI counterstaining (blue) (**supplementary Fig. 3A**).

QuPath was used to randomly sample five 561μm x 561μm areas per region per case. Regions analysed were the medial and lateral substantia nigra, and the medial temporal and occipital striate cortical gray matter. Pericytes were included according to a strict inclusion criteria based on the method of Ding *et al*.^34^ Pericyte cell bodies were identified by their morphology and staining characteristics for quantification. Specifically, the pericyte appeared as a protrusion lining the outer capillary wall and was surrounded by a collagen IV positive ring (membrane) with a clearly labelled DAPI-positive nucleus. Nucleated cell bodies lining the inner capillary wall were not included, as these represent endothelial cells. The number of pericytes on normal capillaries were counted in each of the areas sampled and averaged per case.

#### Quantitation of aquaporin-4 localisation in astrocytic endfeet

AQP4 in perivascular astrocytic endfeet was identified using immunofluorescent sections stained for AQP4 (red) and counterstained with DAPI (blue) to identify the nuclei of endothelial cells lining a blood vessel (**supplementary Fig. 4**). The blood vessels sampled for analysis ranged from approximately 20-60μm in size.

QuPath was used to sample six 561μm x 561μm areas per region per case. Regions analysed were the medial and lateral substantia nigra, and the medial temporal and occipital striate cortical gray matter. The quantitative method of Zeppenfeld *et al*.^35^ was used to determine the amount of AQP4 in astrocytic endfeet in relation to global expression of AQP4 in each of the areas sampled. As we identified equivalent changes in the PVS of both venules and arterioles (see Results), we did not discriminate between these vessel types for the purpose of AQP4 analysis. The fluorescent channel for AQP4 was selected in the digital images of the areas sampled. The mean gray intensity value of the entire area was determined and used as a measure of total AQP4 expression in that sample. DAPI was used as a confirmatory stain to verify the identity of blood vessels and select those meeting the size criteria. A perivascular border of 10μm around each selected blood vessel was drawn and the mean gray intensity value of AQP4 immunofluorescence determined. The relative amount of AQP4 in astrocytic endfeet was calculated as a ratio of perivascular border AQP4 intensity relative to global AQP4 intensity using the following formula:

AQP4 endfeet ratio = perivascular border AQP4 mean gray value / global AQP4 mean gray value

#### Total astrocyte density

Astrocytes were identified using a cocktail of astrocyte markers (details in **supplementary Table 1** for antibody details and **supplementary Table 2** for tissue section stained) and immunoperoxidase detection with a haematoxylin counterstain.

QuPath was used to randomly sample five 561μm x 561μm areas per region per case. Regions analysed were the medial and lateral substantia nigra, and the medial temporal and occipital striate cortical gray matter. Astrocytes in the tissue sections were included according to a strict criteria. Specifically, astrocytes with a clearly defined cell body with a peroxidase-stained cytoplasm surrounding a haematoxylin-stained nucleus, regardless of the absence or presence of processes, were counted. The total number of astrocytes were counted in each of the areas sampled and averaged per case.

#### Statistical analysis

Statistical analyses were performed using GraphPad Prism version 10.2.3 and IBM SPSS Statistics 26. A p-value of <0.05 was identified as the level of significance. Group differences in demographics and pathological severity were assessed using t-tests and Mann-Whitney U tests. Group differences in A, B and CVD scores were assessed using chi-square tests. Multivariate regression analysis was performed to examine structures associated with capillaries, including string vessels, pericytes on capillaries and pericytes on string vessels. Additionally, two separate univariate regression analyses were conducted to model structures related to arterioles and venules (PVS) and astrocytes (AQP4). Age and post-mortem delay were included as covariates in all models. Specific details can be found in the relevant results sections. Spearman rank correlations were also performed to determine whether the measured variables were related to disease duration, A score, B score or LB stage.

## Results

### Demographic and pathological differences between groups

The average age of the PD group was 77 ± 7 years, with an average post-mortem delay of 22 ± 17 h. The average age of the control group was 79 ± 8 years, with an average post-mortem delay of 15 ± 10 h (**Table 1**). There were no significant differences in age at death (t-test *P* > 0.05) or post-mortem delay (t-test *P* >0.05) between the groups.

Consistent with pathological diagnosis, α-synuclein-positive Lewy pathology was only detected in the PD group (**Table 1**). There was no significant difference in A (χ^2^(3) *P* > 0.05), B (χ^2^(2) *P* >0.05) or CVD (χ^2^(1) *P* >0.05) scores between the PD and control group (**Table 1**), indicating an absence of co-existing Alzheimer disease.

### Arteriolar and venular perivascular space size is enlarged in PD

Differences in the venule and arteriole PVS size were analysed in PD and control groups using multivariate regression modelling covarying for age and post-mortem delay, as well as CVD score (presence or absence). There was a significant increase in the average arteriolar PVS size in PD compared to controls (*P* <0.001). Venular PVS size was also significantly increased in the PD group (*P* <0.001). The arteriolar PVS was larger than that of venules, although the ratio of the arteriolar to venular PVS size was not different between PD (mean size ratio= 1.39, *P* > 0.05) and controls (mean size ratio= 1.56, *P* >0.05), indicating both vessel types were affected equally. A summary of the group means and significance can be found in **supplementary Table 3**.

As the arteriolar and venular PVS size were equally affected in PD, we combined the measures for use in future analyses. **Figure 1C** shows the group differences for this combined measure, demonstrating the PVS size in PD is 56% greater than the control value (*P* <0.001).

**Figure 1.**
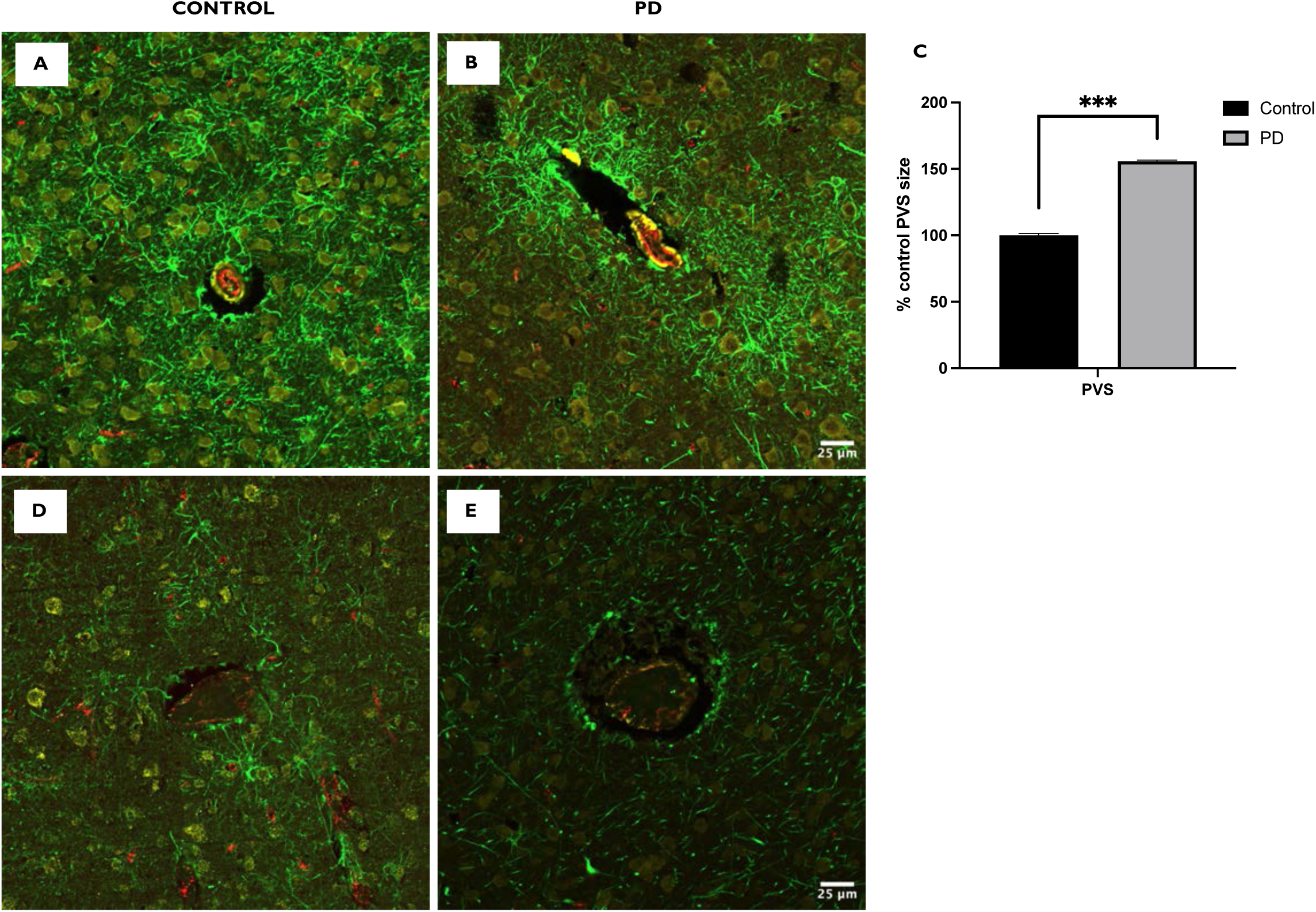
The perivascular space is significantly enlarged in PD compared to controls. Micrographs of sampled arterioles in a representative control **(A)** and PD **(B)** case. Micrographs of sampled venules in a representative control **(D)** and PD **(E)** case. Group differences in PVS size (arterioles and venules combined) graphed **(C)**. All values are expressed as a percentage of the controls. Scale in **(B)** and **(E)** is equivalent for **(A)** and **(D)**. Error bars represent standard error of the mean. *** *P* < 0.001. *n* = 25. PVS = perivascular space.

### The microvasculature is altered in PD

The total length of both capillaries and string vessels were measured in each case (**supplementary Fig. 2**). Differences in normal capillaries and string vessels were analysed using multivariate regression modelling covarying for age and post-mortem delay. There was no difference in normal capillary length in PD and controls (*P* >0.05). The percentage of string vessels was significantly increased in PD compared to controls, representing a 135% increase over control value (**Figure 2A-C**) (*P* <0.001). A summary of the group means and significance can be found in **supplementary Table 3**.

**Figure 2.**
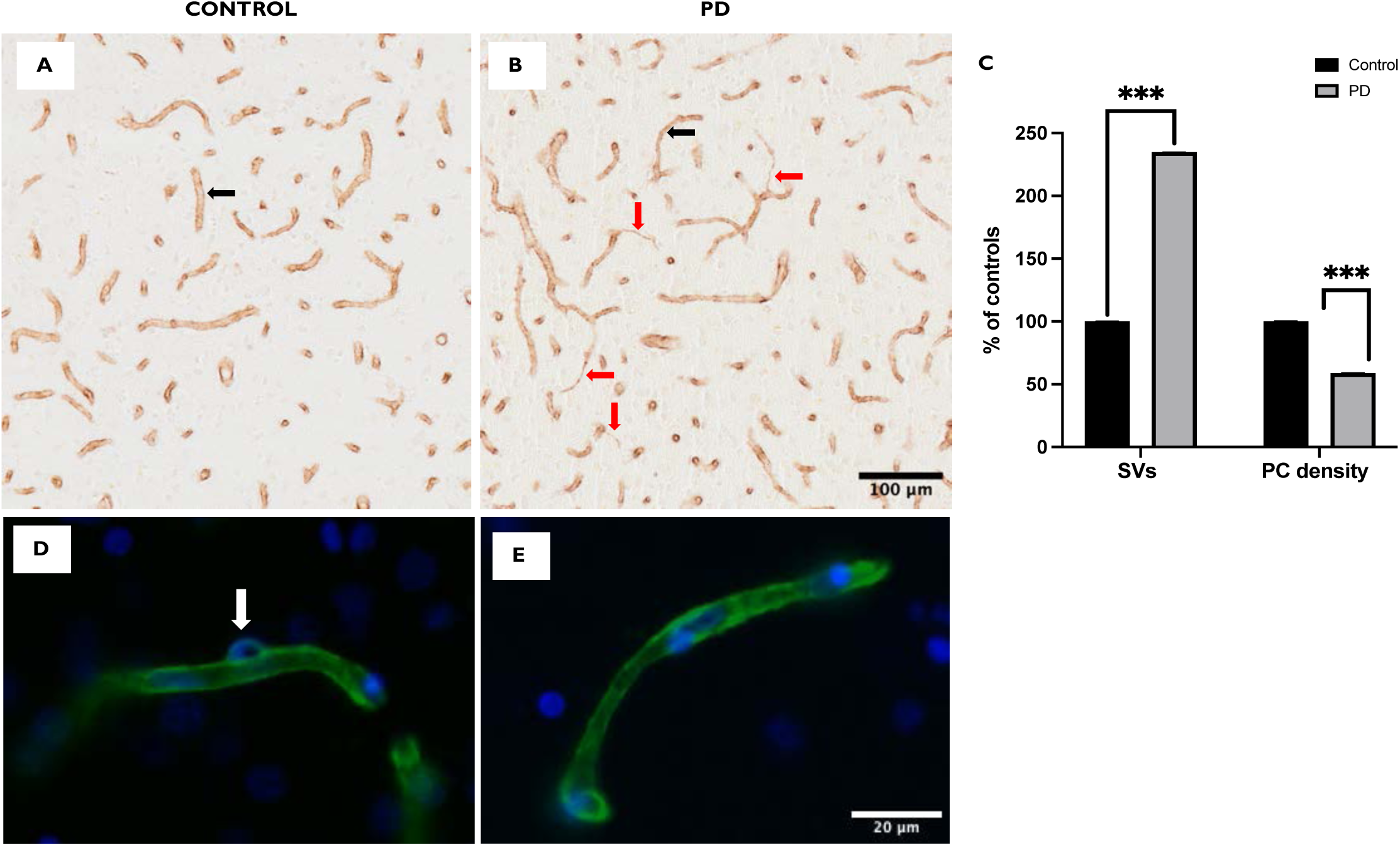
String vessels and pericytes are significantly altered in PD. Micrographs of normal capillaries (black arrows) and string vessels (red arrows) in a representative control **(A)** and PD case **(B)**. Micrograph of pericyte on a normal capillary (white arrow) in a representative control **(D)** and the absence of pericytes in a PD case **(E)** (white arrow). Group differences in the formation of string vessels and density of pericytes is graphed **(C)**. All values are expressed as a percentage of the controls. Scale bar for **(B)** is equivalent for **(A)**. Scale bar for **(E)** is equivalent for **(D)**. Error bars represent standard error of the mean. *** *P* < 0.001. *n* = 25. SV = string vessels. PC = pericytes.

### Cellular regulation of the capillaries is decreased in PD

Pericytes regulate capillary blood flow.^36^ The density of pericytes on capillaries was determined (**supplementary Fig. 3**). Analysis revealed an absence of pericytes on string vessels, consistent with the absence of endothelial cells.^26^ Analysis of pericyte density was therefore quantified on normal capillaries. Differences in capillary-associated pericyte density between the PD and control groups were analysed using multivariate regression modelling for structural measures covarying for age and post-mortem delay. Pericyte density significantly decreased in PD compared to controls, representing 59% of the control value (**Figure 2C-E**) (*P* <0.001). A summary of the group means and significance can be found in **supplementary Table 3**.

### Aquaporin-4 localisation to astrocytic endfeet is decreased in PD

Astrocytic endfeet form a part of the BBB that regulates the transfer of substances in and out of the brain.^37^ AQP4 is the water channel responsible for fluid exchange and homeostasis which is localised to the endfeet of astrocytes.^38^ The total pool of astrocytes was identified using a cocktail of astrocyte markers and their density was quantified. There was no significant difference in astrocyte density between controls and PD patients (univariate analysis, *P* >0.05). The intensity of AQP4 labelling in astrocytic endfeet (AU) was quantified and the ratio of AQP4 localised to astrocytic endfeet versus global levels measured (perivascular endfeet:total AQP4). Differences in AQP4 ratio between the PD and control groups were analysed using univariate regression modelling covarying for age, post-mortem delay, PVS size and total density of astrocytes. AQP4 localisation to astrocytic endfeet was significantly decreased in PD compared to controls, representing 79% of the control value (**Figure 3**). A summary of the group means and significance can be found in **supplementary Table 3**. The differences observed in the cerebrovasculature between PD and the control group (and regional differences) are summarised in **Figure 6**.

**Figure 3.**
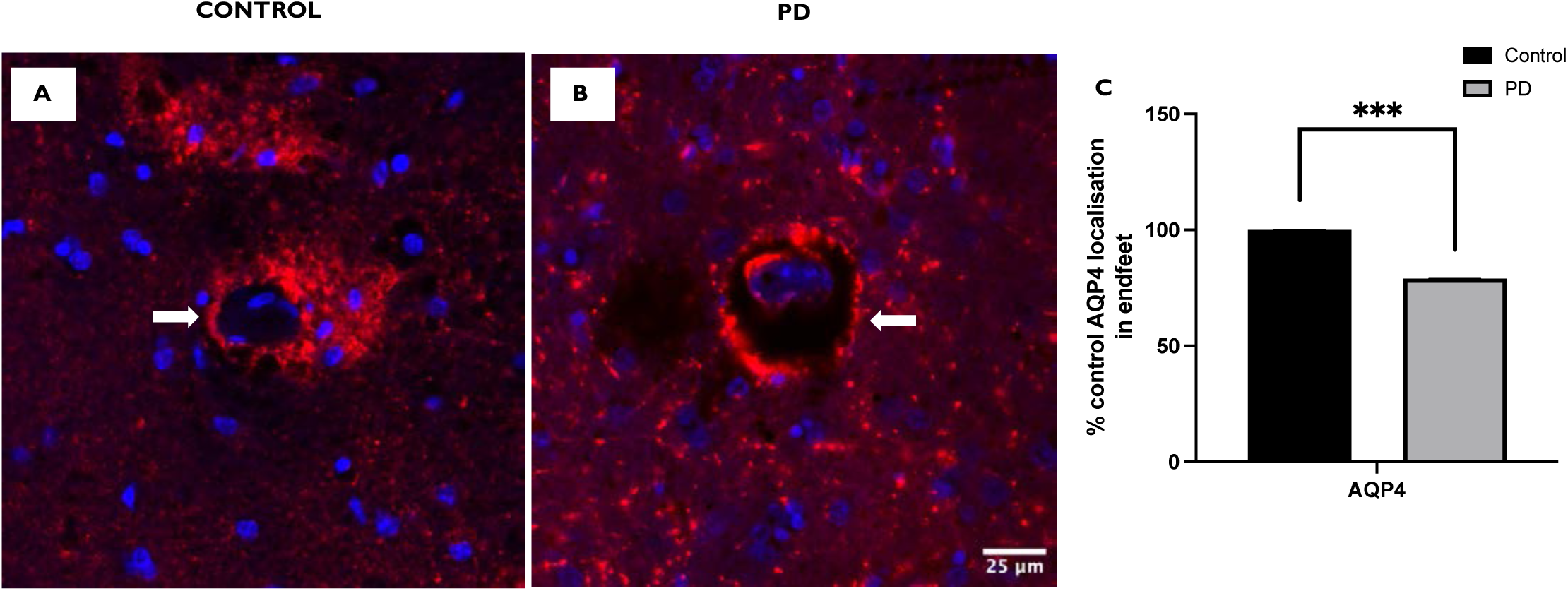
Aquaporin-4 localisation to astrocytic endfeet is decreased in PD compared to the control group. Micrograph of AQP4 in the perivascular border of a blood vessel (white arrow) in a representative control **(A)**. Micrograph showing an obvious reduction of AQP4 from the perivascular border (white arrow) in a PD **(B)** case with the group differences in the perivascular endfeet:total AQP4 ratio graphed **(C)**. All values are expressed as a percentage of the controls. Scale bar for **(B)** is equivalent for **(A)**. Error bars represent standard error of the mean. *** *P* < 0.001. *n* = 25. AQP4 = aquaporin-4.

**Figure 4.**
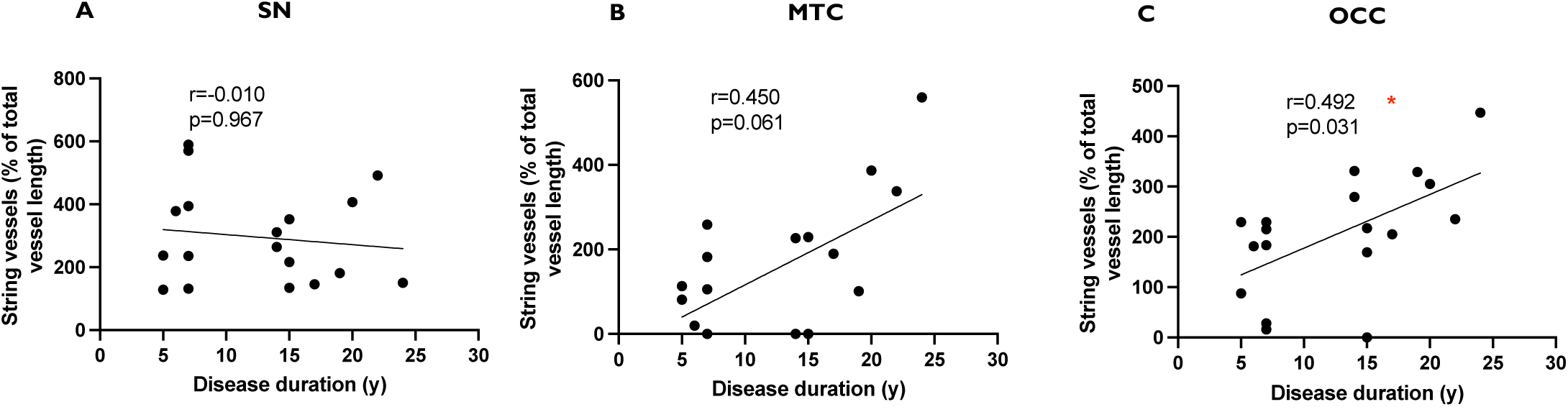
Correlations between the percentage of string vessels and disease duration in variably affected brain regions in PD. Scatter plot demonstrates that the percentage of string vessels was not related to disease duration in the substantia nigra **(A)** or medial temporal cortex **(B)**. Scatter plot demonstrates a significant association between the percentage of string vessels and disease duration in the occipital cortex **(C)**. *Spearman’s Rho *P* <0.05. *n* = 18. SN = substantia nigra, MTC = medial temporal cortex, OCC = occipital cortex.

### Staged regional pathological changes

The regions selected for this study are affected by PD at different stages of pathological progression. The substantia nigra exhibits early pathological deposition of α-synuclein well before cell loss and the onset of clinical motor symptoms,^39^ making it a crucial region for studying early disease-related changes. The medial temporal cortex (part of the limbic system) is affected by pathology during the symptomatic stage of disease and is involved in cognition, while the occipital cortex, though generally unaffected by pathology, is involved in the development of visual hallucinations, which are common in the later stages of PD.^40^ Thus, each region was selected to represent different stages in disease progression.

### Association between protein deposition and regional changes in the cerebrovasculature in PD

To determine if the cerebrovascular changes observed were progressive, Spearman rank correlations were performed for all regional measures in PD cases against pathological protein stages. Although no LB pathology was detected in the occipital cortex, there was a correlation between increasing LB stage and total PVS size in this region (Rho =0.576, *P* <0.05), indicating an increasing enlargement of PVS with pathological disease progression. In contrast, no associations were observed between LB stage and cerebrovascular changes in the medial temporal cortex or substantia nigra.

Further analysis of pericytes on normal capillaries demonstrated regional differences in PD compared to controls (**Figure 5A-E**). In controls, more pericytes were observed on normal capillaries in the occipital cortex compared to the medial temporal cortex or substantia nigra (**supplementary Table 4**). Multivariate regression modelling covarying for age and post-mortem delay found that the density of pericytes on normal capillaries significantly decreased in the occipital cortex in PD, representing 52% of the control value (*P* <0.001). Pericyte density was also significantly decreased in the substantia nigra in PD, reaching 45% of the control value-a greater reduction than observed in the occipital cortex (*P* <0.001). This suggests nigral pericytes are affected early in PD. There was no change in the medial temporal cortex (**Figure 5E**). A summary of the regional means and significance can be found in **supplementary Table 4**.

**Figure 5.**
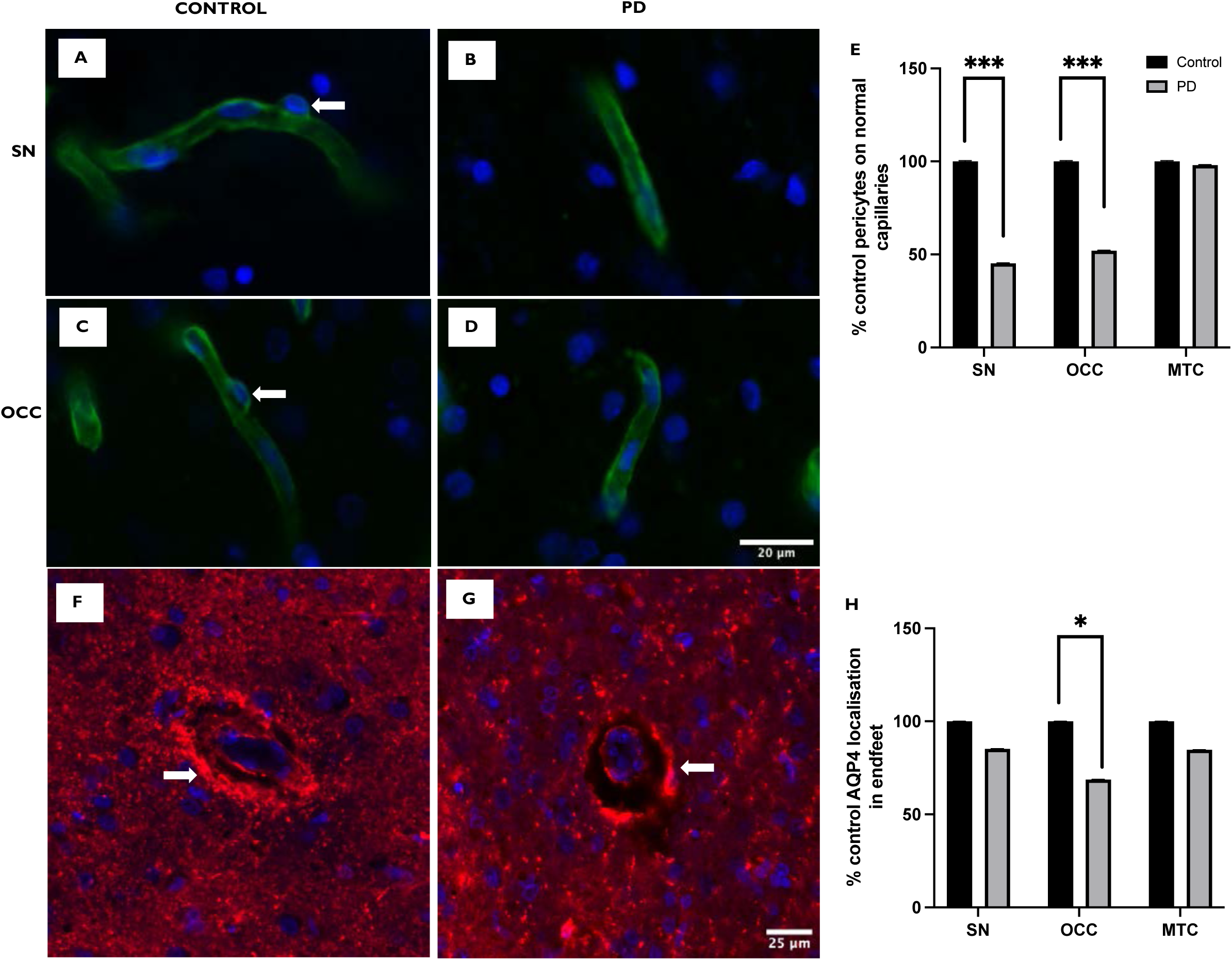
The cerebrovasculature varies in a region-dependent manner in PD compared to controls. Micrographs of pericytes on normal capillaries (white arrow) in representative control cases in the substantia nigra **(A)** and occipital cortex **(C)**. Micrographs showing a reduction of these pericytes in PD cases in the substantia nigra **(B)** and occipital cortex **(D)**. Regional differences in the density of pericytes on normal capillaries between the groups is graphed **(E)**. Micrograph of AQP4 localised to the perivascular border (white arrow) in a representative control case in the occipital cortex **(F)**. A noticeable reduction of AQP4 from the perivascular border (white arrow) can be seen in the PD case **(G)** in the occipital cortex, with the regional differences in the AQP4 perivascular:total ratio between the groups graphed **(H)**. All values are expressed as a percentage of the controls. Scale bar for **(D)** is equivalent for **(A)**, **(B)** and **(C)**. Scale bar for **(G)** is equivalent for **(F)**. Error bars represent standard error of the mean. *** *P* < 0.001, \**P* < 0.05. *n* = 25. PC = pericytes, SV = string vessels, AQP4 = aquaporin-4, SN = substantia nigra, MTC = medial temporal cortex, OCC = occipital cortex.

**Figure 6.**
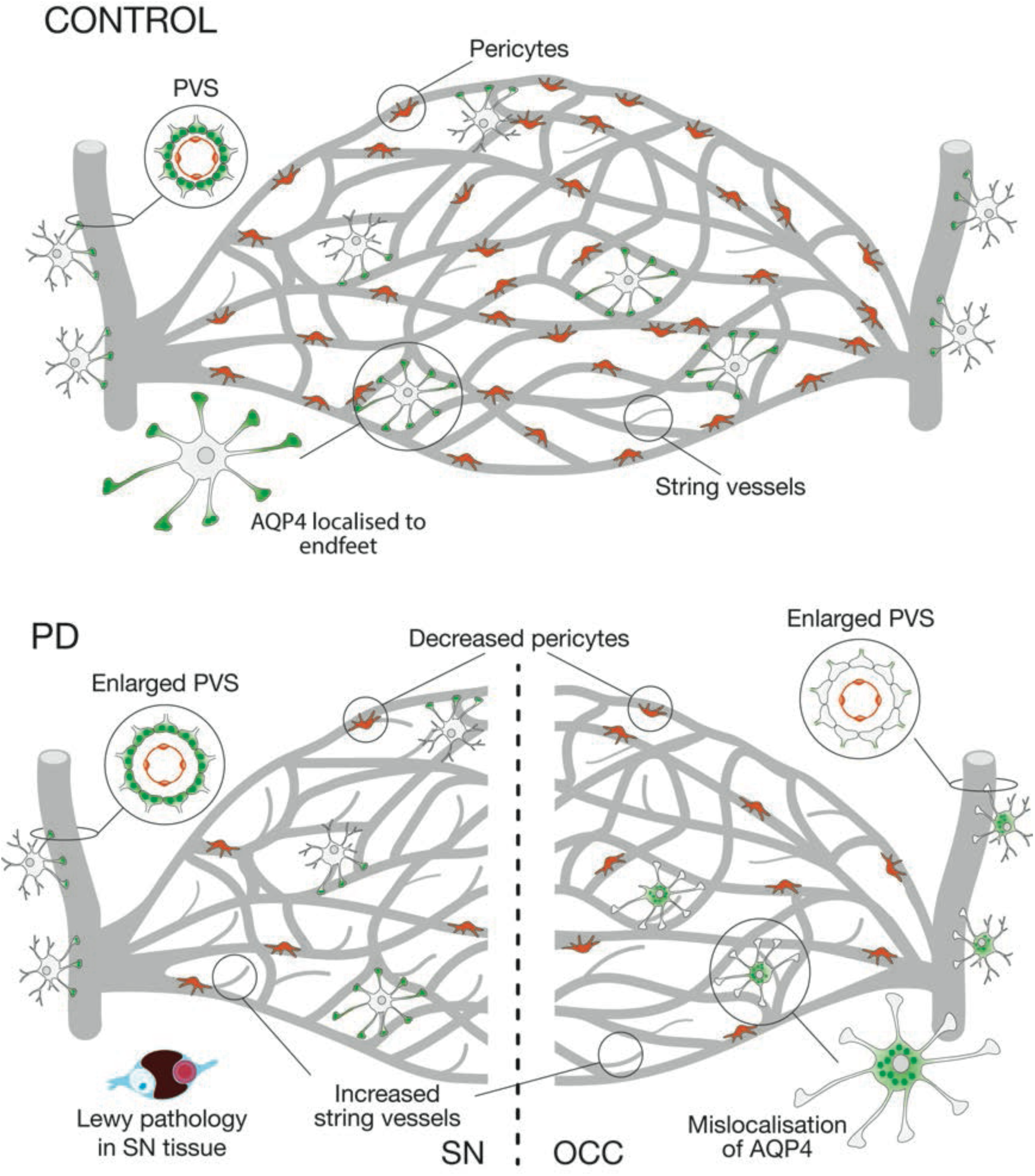
A schematic representing the variations in the cerebrovasculature in PD in the substantia nigra and occipital cortex compared to healthy controls. Compared to controls, the substantia nigra and occipital cortex revealed increased string vessel formation and decreased pericyte density. Enlarged PVS was observed in PD, while mislocalised AQP4 was evident only in the occipital cortex. Lewy pathology was not present in the occipital cortex. PD= Parkinson’s disease, PVS = perivascular space, AQP4 = aquaporin-4, SN = substantia nigra, OCC= occipital cortex.

### Structural vessel changes correlated with disease duration in a regionally staged manner in PD

To determine if the cerebrovascular changes observed were related to disease duration, Spearman rank correlations were performed for all regional outcome measures in PD cases. The analysis identified that disease duration correlated with an increasing proportion of the capillary bed collapsing to string vessels, most marked in the later affected occipital cortex (**Figure 4C**) (Rho = 0.492, *P* <0.05). Capillary string vessel formation in the medial temporal cortex (Rho = 0.450, *P* = >0.05) (**Figure 4B**) and substantia nigra did not correlate with disease duration (Rho = −0.010, *P* = >0.05) (**Figure 4A**). A summary of the regional correlations and significance can be found in **supplementary Table 5**.

### Region-dependent decreases in AQP4 in astrocytic endfeet in PD

While the intensity of AQP4 in astrocytic endfeet was not associated with either pathological progression or disease duration, regional differences in the AQP4 ratio between PD and controls were analysed using univariate regression modelling covarying for age, post-mortem delay, PVS size and total density of astrocytes, as posterior cortical regions are known to have a faster glymphatic clearance rate.^41^ As expected, controls had the highest AQP4 levels in the occipital cortex, then the substantia nigra, and the least in the medial temporal cortex **(supplementary Table 4)**. PD cases had more AQP4 in the substantia nigra than in either cortical region **(supplementary Table 4)** with the occipital cortex bearing the brunt of this relocation of AQP4 (PD representing 69% of the control value, *P* <0.05, **Figure 5F-H**). A summary of the regional means and significance can be found in **supplementary Table 4**.

## Discussion

This study investigated changes in functionally relevant cellular characteristics of the cerebrovasculature in PD, focusing on how these changes relate to disease progression and protein deposition. We compared age- and post-mortem interval-matched control and PD cases with varying disease durations, selecting regions differentially affected by the disease process. Although no significant differences in diagnostic CVD scores were found between groups,^33^ our assessment of structures involved in the glymphatic system showed a notable enlargement of the small arteriolar and venular PVS in PD, along with a decrease in AQP4 from around these vessels. Advancing LB stage was linked to the increased PVS size in the occipital cortex despite no local deposition of α-synuclein in this region, suggesting that such cerebrovascular changes occur prior to significant α-synuclein (or other protein) deposition. In addition to these glymphatic changes, significant alterations in the capillary network that impact blood flow and its regulation were observed, including an increase in string vessel formation and a reduction in pericyte density. Correlative analyses showed that capillary string vessel formation in the occipital cortex increased with disease duration. While greater pericyte loss was observed in the substantia nigra where significant neuronal loss occurs, the medial temporal cortex remained unaffected despite significant α-synuclein deposition. In the regions analysed with α-synuclein deposition, there were no associations between vascular changes and progressive pathological severity. These findings underscore that progressive small vessel cerebrovascular changes in PD largely occur independently of α-synuclein deposition. Regionally, the occipital cortex appears particularly susceptible to these progressive cerebrovascular disease changes during the disease process.

### Progressive changes in the perivascular space and BBB integrity occur in PD

Cerebrovascular disease has been extensively studied in Alzheimer’s disease,^42–45^ but research on its role in Lewy body disease is sparse and even more so in PD. In Alzheimer’s disease, protein deposition is known to compromise the cerebral vasculature and BBB integrity,^46,47^ but this is not the case in PD. In the current study, cerebrovascular changes occurred in the absence of α-synuclein pathology in the visual cortex and were limited in the medial temporal cortex where significant α-synuclein pathology occurs. This suggests that the cerebrovascular changes occur independently of α-synuclein deposition. A systemic review of blood flow changes in PD^48^ identifies regions with limited LB deposition^4^ as consistently having reduced blood flow in PD, including the occipital lobe and visual network, as well as in the caudate/putamen, supplementary and sensory networks. These later regions correlate with PD related motor dysfunction, and with regional reductions in metabolism (PET glucose) and function (fMRI).^48^ A similar regional pattern of increased cerebrovascular reactivity latency has been recently observed,^49^ supporting the concept that functional cerebrovascular changes impact brain regions with limited α-synuclein deposition. Our data suggest that this is due to regional structural changes in cerebral vasculature in PD.

Although not all cerebrovascular changes observed through neuroimaging, such as white matter hyperintensities and cerebral microbleeds, can be confirmed neuropathologically,^50^ advanced MRI resolution has proven effective in reliably detecting PVS size.^51,52^ Brain imaging studies have identified dilation of the PVS in the midbrain,^53,54^ caudate/putamen,^52,54,55^ and in the centrum semiovale^54,55^ in PD. Additionally, human brain tissue studies have confirmed enlargement of the PVS in nigrostriatal regions, including the brainstem and caudate/putamen,^56^ consistent with our current observations. Until recently, enlarged PVS were considered largely benign, although studies have now identified associations with motor deficits,^57^ and cognitive decline in PD,^58,59^ highlighting a potential role in disease progression.

The mechanism behind enlarged PVS remains elusive, although it has been suggested that such changes may be reflective of BBB dysfunction.^52^ It is widely established that compromised integrity of the BBB can lead to heightened vascular permeability and subsequent widening of the PVS due to excess fluid accumulation.^60,61^ This accumulation of fluid is thought to arise from impaired or obstructed drainage. Aquaporins (AQP) are transmembrane water channel proteins localised to the endfeet of astrocytes in the brain which regulate water diffusion intra- and intercellularly.^62^ Together, there are 13 known aquaporin isoforms, with AQP1, AQP4 and AQP9 all notable in the CNS.^63^ AQP4 is the most abundant water channel in the human brain^64^ and facilitates bi-directional movement of water between the blood vessels, the astrocyte cytoplasm and the parenchyma (perivascular CSF-interstitial fluid exchange).^65,66^ Complete dysfunction or decrease of AQP4 in astrocytic endfeet disrupts the fluid exchange, ultimately resulting in abnormal interstitial fluid accumulation.^67,68^ In aging mice, impaired CSF recirculation and interstitial fluid clearance has been demonstrated with a decrease in AQP4 around enlarged PVS size,^69^ supporting a role for AQP4 in PVS enlargement.

Impaired fluid drainage of the brain through glymphatic system dysfunction has been identified in PD. Diffusion tensor MRI measures water diffusion along white matter tracts and is specifically used to measure water diffusion along the perivascular space (DTI-ALPS) as a measure of the glymphatic flow.^70^ Recent studies show progressive reductions in DTI-ALPS in PD patients with Hoehn and Yahr stages >2 compared to healthy controls,^57,71–74^ associated with increasing disease severity^71,73^, disease duration,^57,75^ levodopa equivalent daily dose,^75^ and cognitive impairment.^71,75^ Reductions in DTI-ALPS correlates with enlarged PVS size,^73,76^ with both measurements reflective of glymphatic dysfunction in PD. Fluid drainage from the brain is via the meningeal lymphatic vessels, and with increasing PD stage, uptake of fluid into this drainage system is significantly impaired without any meningeal structural changes.^77^ This is consistent with a retention of parenchymal fluid in PD. The positioning of the occipital cortex close to the posterior meningeal drainage system may be of significance for the changes we have observed, as the greatest fluid drainage into the meningeal lymphatics in people with mild stroke is posteriorly and directly related to enlarged PVS size.^78^ Interestingly, in mice injected with α-synuclein preformed fibrils, occlusion of the meningeal lymphatics^77^ or a reduction in astrocytic AQP4^79^ increases α-synuclein pathology and exacerbates motor and memory deficits. In our study, we show that increasing LB stage is associated with increasing PVS size but not reduced AQP4 in astrocytic endfeet. This is consistent with both increasing PVS and more widespread LB formation contributing to the progression of PD.

Our findings support a previous study demonstrating reduced localisation of AQP4 to astrocytic endfeet in PD.^24^ Indeed, Braun *et al*.^24^ reported a reduction in AQP4 localisation in the frontal cortex of human post-mortem cases with neocortical Lewy body pathology. This change was accompanied by α-synuclein accumulation in the form of Lewy bodies, suggesting that the loss of perivascular AQP4 may relate to the increase in pathology in this region.^24,80^ As stated above, our study confirms reduced AQP4 in astrocytic endfeet in PD, even in regions without α-synuclein pathology, with only enlarged PVS relating to increasing LB stage. If parenchymal fluid retention occurs prior to PVS enlargement, the changes in astrocytic AQP4 and/or changes in pulsatile activity are likely to contribute to the PVS enlargement. Recent studies show that intracranial pressure pulsatility that is required for parenchymal fluid flow is directly linked to heartbeat and breathing rhythms through neural mechanosensitive channels that modulate the brains electrical rhythmicity.^81,82^ Patients with PD have a 2.6 fold longer QT interval on their ECG than controls, and this decrease in heartbeat is associated with a greater severity of PD and a greater probability of developing more severe PD.^83^ Reduced noradrenergic innervation of the heart assessed using metaiodobenzylguanidine (MIBG) correlates with occipital hypoperfusion in PD,^84^ suggesting these factors are related.

### Regional degeneration of the capillary network occurs in PD

Alongside the cerebrovascular changes observed in larger calibre vessels, our results also revealed significant capillary degeneration in PD, marked by a notable increase in string vessel formation and a decrease in pericyte density. Our findings of increased string vessel formation align with previous studies of vascular degeneration in PD that have demonstrated morphological changes at the capillary level of the cerebrovasculature.^25,26,28^ Specifically, Yang *et al*.^26^ observed a significant increase in string vessels in the substantia nigra, which is thought to be a consequence of endothelial cell degeneration.^25^ Consistent with this degenerative hypothesis, we observed an increase in string vessels and decline in pericytes on normal capillaries in PD compared with controls. String vessels have been linked to cerebral hypoperfusion,^85,86^ with hypoperfusion common in the occipital and visual network, as well as in the caudate/putamen, supplementary motor and sensory motor networks in PD, as described above. Hypoperfusion has been shown to promote endothelial cell death, ultimately causing the collapse of the capillaries into string vessels, thereby driving a cycle of degeneration.^85–87^

Our results also demonstrated a significant loss of capillary-associated pericytes in PD, with changes seen even on normal capillaries. Pericytes are mural cells situated on the abluminal surface of capillaries and are essential for preserving the integrity of the BBB, as well as the regulation of cerebral blood flow.^88^ As contractile cells, pericytes can control blood flow.^36^ Reduced pericyte number has previously been reported in the grey matter of the middle frontal gyrus in PD.^28^ This reduction was observed in regions where endothelial degeneration was present.^25^ This indicates that the loss of pericytes may compromise the integrity of the capillary walls, as studies have shown severe vascular damage in pericyte deficient mouse models.^89^ Moreover, it is difficult to separate pericyte loss and string vessel formation, given the intricate cross-talk between pericytes and endothelial cells.^28,89,90^ Indeed, Bell *et al*.^89^ observed a reduction in tight junction protein expression by endothelial cells in pericyte-deficient mouse brains. This loss in tight junctions heightens the permeability of the BBB and causes degeneration of endothelial cells.^89,91^ In the present study, the reduction in pericytes was observed on normal capillaries, suggesting that degeneration of pericytes occurs prior to string vessel formation. This is consistent with previous studies showing that pericyte loss affects capillary flow and structure during aging,^92^ which is accelerated in ApoE4 carriers leading to endothelial cell degeneration.^93^ Animal models have also shown that pericyte loss can lead to BBB breakdown, reduced cerebral blood flow and accelerated neurodegeneration.^89,94,95^ Although our findings did not reveal a direct, regional correlation between pericyte loss and the presence of α-synuclein, mouse models of PD have shown that an over-expression of α-synuclein leads to pathological activation of pericytes at an early stage, even prior to changes in pericyte density,^96^ suggesting that they may react first to remote α-synuclein pathology.

### Sparing of the cerebrovasculature in the medial temporal cortex in PD

Our observations revealed regional variability in cerebrovasculature alterations in PD with marked sparing of the medial temporal cortex. One explanation for this finding may be due to regional variations in vascular innervation, BBB integrity and blood flow to this region. In *de novo* PD patients, the hippocampus is relatively hyperperfused compared with the hypoperfusion observed in the occipital lobe,^97^ a finding that continues to be observed into advanced disease.^98^ Limited reductions in blood flow in the hippocampus is a feature of the PD-related patterns discussed earlier. Hippocampal arterioles have been shown to be functionally unique due to their organisation and the neurochemistry of their endothelium,^99^ potentially contributing to their protection in PD. Proteomic analysis of the hippocampus in human post-mortem PD tissue has also revealed an upregulation of α-1-syntrophin in this region, which localises AQP4 to astrocytic endfeet,^100^ consistent with our findings of normal AQP4 levels in astrocytic endfeet in this region. Lewy pathology in the hippocampus is found in the CA2/3 subfields^4,101^ without evident cerebrovascular pathologies. Further work to determine potentially modifiable factors that protect the hippocampus from the small vessel cerebrovascular changes observed elsewhere in PD may provide novel treatment options.

### Study limitations and considerations

There are methodological considerations that should be accounted for when interpreting the above results. Firstly, it is important to note that we only assessed the presence of insoluble α-synuclein, β-amyloid and tau, and found no direct association between the deposition of these pathological proteins and the vascular changes observed. Previous animal studies have identified alterations in soluble α-synuclein,^24^ which are unlikely to be detectable by immunohistochemical analysis of formalin fixed paraffin-embedded tissue sections. This warrants further investigation into changes in protein deposition using proteomic techniques, such as western blot. We acknowledge that the sample size of our control group was small, as we aimed to include only age matched, disease-free cases. This was determined by availability at the Sydney Brain Bank, as we wanted to avoid introducing samples from different brain banks and potential case discrepancies due to variations in sample handling and preparation, such as fixation, which can impact study outcomes. Further, we identified pericytes in our tissue sections based on strict morphological criteria.^34^ Although our observations of pericyte loss in PD aligned with previous findings,^28^ the availability of a reliable pericyte marker (specific to just pericytes) would have been useful in terms of identifying biochemical changes, and whether those changes occur prior to the degeneration or loss of this cell type.^34^

## Conclusion

In conclusion, our findings revealed significant regional changes in the PVS, capillaries and pericytes in PD, along with changes in AQP4. These changes are progressive and occur in regions where α-synuclein deposition is more limited, although animal models of PD show that changes in fluid dynamics increase α-synuclein accumulation. While the most affected brain regions have previously been considered unaffected by the disease process, our findings highlight significant cellular changes that appear to align with hypoperfusion, consistent with the type of vascular pathologies observed. These results suggest that targeting these changes could provide therapeutic benefits for mitigating the cellular progression of PD.

## Data availability

The data generated for this study is available upon request from the corresponding author.

## Acknowledgements

Tissues were received from the Sydney Brain Bank which is supported by Neuroscience Research Australia and is a special gift in memory of Jim Raftos from the Shaw family. We would like to thank Dr Michael Carnell, Dr Sandra Fok and Dr Maria Kasherman from the Katharina Gaus Light Microscopy Facility at the University of New South Wales for assistance with the imaging and analysis. We would like to thank Heidi Cartwright for assistance with the figurework.

## Funding

The research reported in this publication is supported by the Michael and Elizabeth Gilbert Postgraduate Award in Parkinson’s Disease Research awarded to DD, and funding to GH from the National Health and Medical Research Council of Australia [Investigator Grant 1176607].

## Competing interests

The authors report no competing interests.

## Supplementary material

Supplementary material is available at *Brain* online.

## Abbreviations

AQP: aquaporin
AQP4: aquaporin-4
BBB: blood-brain barrier
C: control
cAMP: cyclic adenosine monophosphate
CVD: cerebrovascular disease
DAB: 3,3’-diaminobenzidine
DTI-ALPS: diffusion tensor image analysis along the perivascular space
ISF: interstitial fluid
LB: Lewy body
LN: Lewy neurite
MTC: medial temporal cortex
NFT: neurofibrillary tangle
OCC: occipital cortex
PC: pericyte
PD: Parkinson’s disease
PVS: perivascular space
SN: substantia nigra
SV: string vessel

## Notes

### Competing Interest Statement

The authors have declared no competing interest.

