## Supplementary material for "Region-specific variations in the cerebrovasculature underlie disease progression in Parkinson’s disease"

SUPPLEMENTARY TABLES AND FIGURES

### Supplementary Table 1 Antibodies and reagents used in study

| Antibody/Reagent | Species | Target | Concentration for IHC | Company (Catalog #) |
|---|---|---|---|---|
| Collagen IV | Goat | Basement membrane | 1:500 | Novus Biologicals (NBP1-26549) |
| GFAP | Rabbit | Astrocytes | 1:500 | Dako (Z0334) |
| S100β | Mouse | Astrocytes | 1:250 | Life Technologies (MA5-15359) |
| ALDH1L1 | Mouse | Astrocytes | 1:500 | Origene (UM500040) |
| GFAP | Goat | Astrocytes | 1:200 | My Bio Source (MBS241916) |
| SMA-Cy3 | Mouse | Smooth muscle | 1:10000 | Sigma-Aldrich (C6198) |
| Aquaporin-4 | Rabbit | Aquaporin-4 | 1:100 | Novus Biologicals (NBP1-87679) |
| Factor VIII/Von Willebrand Factor | Rabbit | Endothelial cells | 1:10000 | Dako (A008202) |
| Phospho-tau (AT8) | Mouse | Neurofibrillary tangles | 1:1500 | Invitrogen (MN1020) |
| β-amyloid | Mouse | Amyloid plaques | 1:200 | Dako (M0872) |
| α-synuclein | Mouse | Lewy bodies and Lewy neurites | 1:7000 | BD Biosciences (610787) |
| Anti-goat Cy2 secondary antibody | Donkey | | 1:200 | Jackson ImmunoResearch (705-225-147) |
| Anti-goat Alexa 488 secondary antibody | Donkey | | 1:200 | Invitrogen (A-10037) |
| Anti-rabbit Alexa 647 secondary antibody | Donkey | | 1:200 | Invitrogen (A-31573) |
| Anti-goat biotinylated secondary antibody | Horse | | 1:200 | Vector Laboratories (BA9500) |
| Anti-mouse biotinylated secondary antibody | Horse | | 1:200 | Vector Laboratories (BA2000) |
| Anti-rabbit biotinylated secondary antibody | Horse | | 1:200 | Vector laboratories (BA1100) |
| VECTASTAIN Elite ABC Peroxidase Kit | Universal | | 1:500 | Vector Laboratories (PK6100) |
| 3,3'-diaminobenzidine (DAB) | Universal | | 1:33 | Vector Laboratories (SK4105) |
| TrueBlack Lipofuscin Autofluorescence Quencher | Universal | | 1:20 | Biotium (23007) |
| DAPI and Hoechst Nucleic Acid Stains | Universal | | 1:1000 | Invitrogen (D1306) |

**Supplementary Table 2 Summary of serial sequence and corresponding staining analysis**

| Serial section | Staining analysis performed | Antibodies | Immunohistochemical technique |
|---|---|---|---|
| 1 | Single label α-synuclein | α-synuclein | Peroxidase |
| 3 | Triple label astrocyte stain | GFAP, S100β, ALDH1L1 | Peroxidase |
| 5 | Triple label PVS stain | GFAP, SMA-Cy3, Factor VIII (+DAPI counterstain) | Immunofluorescence |
| 6 | Single label AQP4 stain | AQP4 (+ DAPI counterstain) | Immunofluorescence |
| 7 | Single label string vessel stain | Collagen IV | Peroxidase |
| 8 | Single label pericyte stain | Collagen IV (+ DAPI counterstain) | Immunofluorescence |

**Supplementary Table 3 Group means and significance for the tested variables**

| Variable tested | Mean PD | Mean Control | *P*-value |
|---|---|---|---|
| Arteriolar PVS (μm) | 20.3 ± 0.6 | 13.6 ± 1.0 | *<0.001* |
| Venular PVS (μm) | 14.6 ± 0.6 | 8.7 ± 0.9 | *<0.001* |
| Total PVS (μm) | 34.8 ± 0.8 | 22.4 ± 1.4 | *<0.001* |
| Length of normal capillaries (μm) | 2035.8 ± 57.6 | 2131.0 ± 95.5 | >0.05 |
| Percentage of string vessels (%/μm) | 1.1 ± 0.1 | 0.5 ± 0.1 | *<0.001* |
| Pericyte density on normal capillaries (/mm$^2$) | 5.1 ± 0.2 | 8.6 ± 0.3 | *<0.001* |
| Total astrocyte density (/mm$^2$) | 31.4 ± 0.5 | 29.4 ± 0.9 | >0.05 |
| AQP4 perivascular endfeet:total ratio (AU) | 1.1 ± 0.3 | 1.4 ± 0.1 | *<0.001* |

Between group comparisons were assessed by univariate and multivariate regression analyses
Values are presented as mean ± SE
PVS perivascular space
AQP4 aquaporin-4
AU arbitrary unit

**Supplementary Table 4 Regional means and significance for the tested variables**

| Region | Variable tested | Mean PD | Mean Control | *P*-value |
|---|---|---|---|---|
| SN | Pericyte density on normal capillaries (/mm$^2$) | 3.6 ± 0.3 | 8.1 ± 0.4 | *<0.001* |
| MTC | Pericyte density on normal capillaries (/mm$^2$) | 5.0 ± 0.3 | 5.1 ± 0.5 | >0.05 |
| OCC | Pericyte density on normal capillaries (/mm$^2$) | 6.6 ± 0.3 | 12.7 ± 0.4 | *<0.001* |
| SN | AQP4 perivascular endfeet:total ratio (AU) | 1.2 ± 0.1 | 1.4 ± 0.1 | >0.05 |
| MTC | AQP4 perivascular endfeet:total ratio (AU) | 1.1 ± 0.1 | 1.2 ± 0.1 | >0.05 |
| OCC | AQP4 perivascular endfeet:total ratio (AU) | 1.1 ± 0.0 | 1.6 ± 0.1 | *<0.05* |

Between group comparisons were assessed by univariate and multivariate regression analyses
Values are presented as mean ± SE
SN substantia nigra
MTC medial temporal cortex
OCC occipital cortex
AQP4 aquaporin-4

**Supplementary Table 5 Regional correlations and significance for tested variables**

| Region | Variable tested | Spearman's Rho | *P*-value |
|---|---|---|---|
| SN | String vessel correlation with DD | -0.01 | >0.05 |
| MTC | String vessel correlation with DD | 0.45 | >0.05 |
| OCC | String vessel correlation with DD | 0.49 | *<0.05* |
| OCC | LB stage correlation with total PVS | 0.58 | *<0.05* |

Spearman's rank correlations were performed
SN substantia nigra
MTC medial temporal cortex
OCC occipital cortex
DD disease duration
LB Lewy body

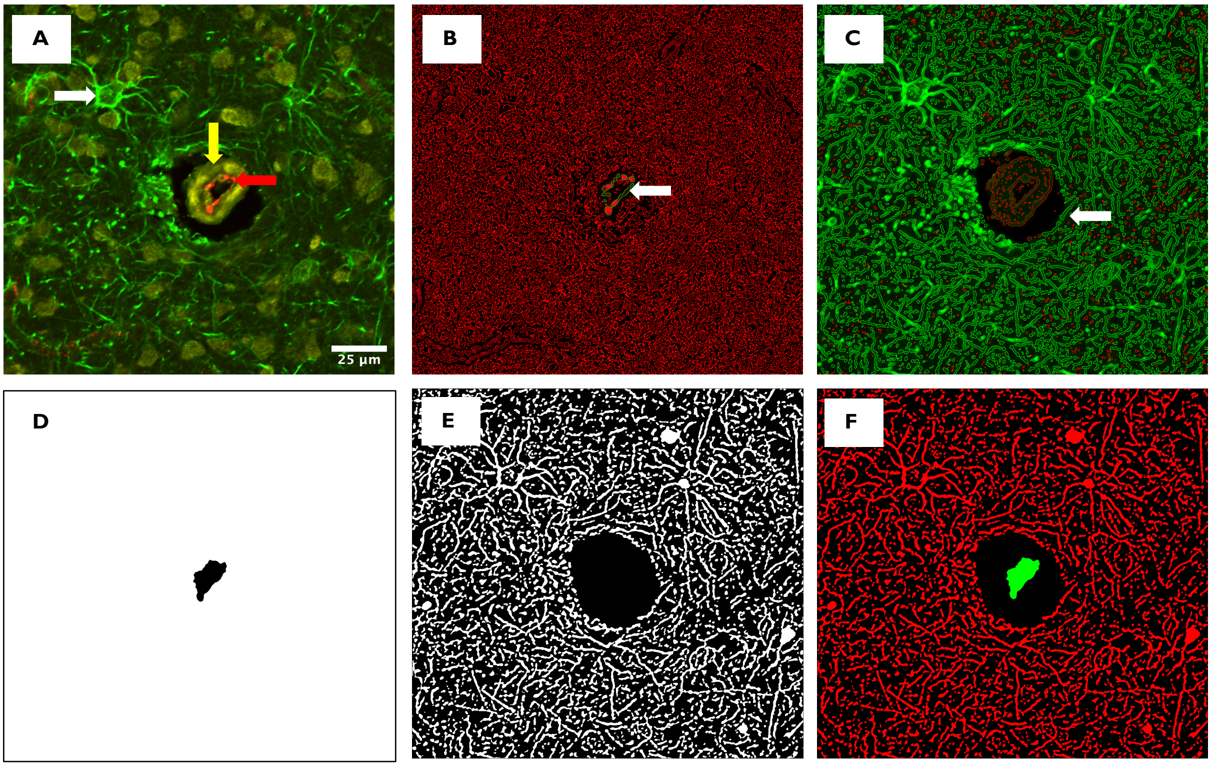


**Supplementary Figure 1 Micrographs showing the workflow for PVS quantitation. (A)** Tissue section stained with GFAP (green; astrocytes, white arrow), SMA (yellow; smooth muscle, yellow arrow) and factor VIII (red; endothelium, red arrow) to identify blood vessels and the perivascular space. The endothelium **(B**; white arrow**)** and outer perivascular wall **(C**; white arrow**)** were identified as objects of interest through pixel intensity. Mask-like images were created to identify inner **(D)** and outer **(E)** boundaries of the blood vessel. **(F)** Both masks were merged and the mean distance (µm) between the two objects, sampled repeatedly to obtain a representative distance, was determined. Please note that the object colours in micrographs **(B)**-**(F)** do not correspond to the fluorophores shown in **(A)**.

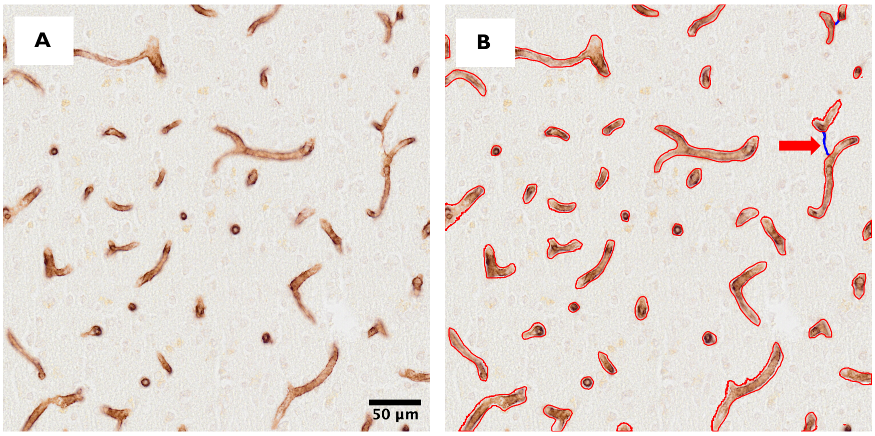


**Supplementary Figure 2 Micrographs showing immunoperoxidase labelling for the analysis of normal capillaries and string vessels.** 5 regions of interest were randomly sampled **(A)** and QuPath was used to recognise capillaries from background tissue (red outline). String vessels (blue outline) were identified by their string-like appearance and selected manually **(B**; red arrow**)**. Scale bar for **(A)** is equivalent for **(B)**.

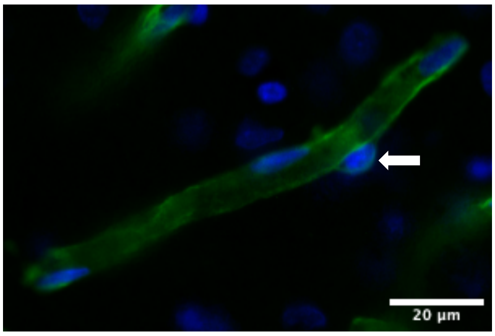


**Supplementary Figure 3 Micrographs showing immunofluorescence labelling for the analysis of pericytes on normal capillaries**. Pericytes were identified by their morphology. The pericyte nucleus was stained with DAPI and appeared as a protrusion (white arrows) on the outer capillary wall that was surrounded by a collagen IV positive membrane.

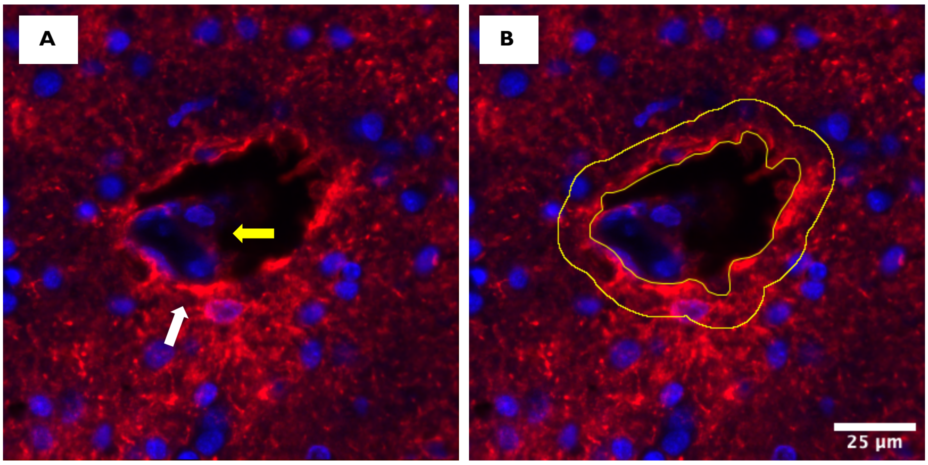


**Supplementary Figure 4 Micrographs showing AQP4 immunofluorescence labelling in perivascular astrocytes.** To assess AQP4 localisation to astrocytic endfeet (white arrow) **(A)**, 6 blood vessels were randomly sampled in the regions of interest, and DAPI was used as a confirmatory stain to ensure structures were indeed blood vessels (yellow arrow) **(A)**. 10µm selection bands were created around the perivascular border and the amount of AQP4 in astrocytic endfeet was calculated as a ratio of perivascular border (endfeet) intensity relative to global, or total AQP4 intensity (perivascular endfeet:total) **(B)**. Scale bar for **(B)** is equivalent for **(A)**.
